# Adapting the ELEAT (Early Life Exposure Assessment Tool) to Portugal – a pilot study to tackle gene-environment interactions in Autism Spectrum Disorder

**DOI:** 10.1101/520593

**Authors:** Celia Rasga, João Xavier Santos, Ana Leonie Lopes, Ana Rita Marques, Joana Vilela, Muhammad Asif, Guiomar Oliveira, Deborah Bennett, Cheryl Walker, Rebecca J. Schmidt, Astrid Moura Vicente

## Abstract

**Background:** Autism Spectrum Disorder (ASD) is a pervasive and clinically heterogeneous neurodevelopmental disorder characterized by deficits in social communication and interaction skills, and repetitive and stereotyped behaviours. It is known that ASD has a strong genetics component, but heritability estimates of 50-80% suggest that modifiable non-genetic factors may play an important role in the onset of the disorder. Recently, pre-, peri and post-natal exposure to a variety of environmental factors has been implicated in ASD. Yet, the comprehensive assessment of environmental exposures in this pathology, using large population datasets, is still lacking. The objective of this study was to pilot an environmental exposure assessment tool in Portugal.

**Methods:** To examine environmental exposures in a population of Portuguese children with ASD, we translated, adapted and piloted the Early Life Exposure Assessment Tool (ELEAT). The ELEAT was originally developed to assess environmental factors in studies of neurodevelopmental disorders. It is a questionnaire filled by mothers of children with ASD, enquiring about Demographic Information, Maternal Conditions/Medical Interventions, Breastfeeding and Child Diet, Maternal Diet, Supplements, Lifestyle, Home and Environment, Environment, Occupation and Exposures. The ELEAT gathers information about environmental exposure along key phases for early neurodevelopment, from 3 months prior to conception, pregnancy, labor and delivery to the first year of life of the child. Two focus groups were realized, one with mothers of typically-developing children and another with mothers of children with ASD, in order to discuss the mothers opinion regarding the tool comprehensiveness and relevance.

**Results:** The large majority of mothers were sure about their answers for all modules, with a small fraction of the group reporting difficulties for the Occupations/Exposures module. Most mothers considered the ELEAT to be a little too long, but generally found that the instructions were clear and, most importantly, agreed that the questions were important.

**Conclusions:** Integration of the pilot feedback will allow us to enhance the tool and optimize its usage in Portuguese-speaking communities, improving its capacity to assemble accurate environmental data from diverse cultural settings, and to be extended to larger population datasets. Combined with genetic and clinical data, the ELEAT will contribute to the identification of modifiable lifestyle and environmental risk factors for ASD. Such evidence may eventually provide the opportunity for disease prevention or reduced severity by mitigating exposure when genetic susceptibility is identified early in life.

## INTRODUCTION

Autism spectrum disorder (ASD) is a complex neurodevelopmental disorder that manifests in early childhood. It is characterized by impairments in communication and social behavior and by repetitive behaviors (American Psychiatric Association, 2013). ASD encloses a group of heterogeneous neurodevelopmental symptoms and occurs in all social, racial, ethnic and socioeconomic groups, with a prevalence of about 9.2 cases per 10 000 children in Portugal (Oliveira et al. 2017. In terms of global prevalence, studies estimate a median ASD prevalence of 17/10 000, a value that varies widely among countries (being smaller in Europe when compared to USA) (Elsabbagh et al., 2012), and that has been markedly rising over the past 30 years (Lai, Lombardo, & Baron-Cohen, 2014; Elsabbagh et al., 2012). There has been much discussion about the nature of this increase, and recent research paints a picture of apprehensiveness. But what makes it so difficult to understand these prevalence increase? Why do we keep searching for alternative explanations?

In the last few decades, autism research has grown exponentially. However, some fundamental questions remain unanswered, namely the ones regarding its causal factors. The etiology of ASD is complex and poorly understood due to the broad variability or clinical heterogeneity of the spectrum. ASD has been demonstrated to be highly modulated by genetic factors (Sandin, Lichtenstein, Kuja-Halkola, Hultman,Larsson & Reichenberg, 2017), with heritability values ranging between 50% and 80%. Nowadays, it is suggested that 10% to 15% of ASD cases are due to multiple rare genetic variants found in patients with the disorder or appear in the context of monogenic syndromes (Abrahams & Geschwind, 2010; Ozonoff, Young, Carter, et al., 2011). However, these instances explain only a minority of cases, with the others remaining idiopathic (Devlin & Scherer, 2012; Zafeiriou, Ververi, Dafoulis, Kalyva & Vargiami, 2013).

Thus, it seems clear that there is more to the story and that ASD onset cannot be solely accounted by genetics. That is, although genetic factors are clearly important, and might be largely responsible for the occurrence of autism, they cannot fully explain all cases. Therefore, it is likely that, in addition to a certain combination of autism-related genes, specific environmental modulators might act as risk factors, triggering the development of autism by acting in concert with genetic susceptibilities (Hallmayer, Cleveland, Torres, Phillips, Cohen, Torigoe, … & Lotspeich, 2011; Grabrucker, 2013).

Given the notion that genetic factors cannot fully explain ASD onset, a considerable number of studies are now focusing on the role of environmental factors in the disorder, an area that was previously underexplored. There is now mounting evidence pointing towards a role played by environmental factors in ASD risk. The developing human brain is understood to be highly susceptible to injury caused by toxic chemicals in the environment. This vulnerability is greatest during embryonic and fetal life, and may be especially important in the first trimester of pregnancy. There are windows of susceptibility in early development that have no counterpart in the mature brain (Rodier, 1995; Rice & Barone, 2000). Furthermore, when mechanisms responsible for body homeostasis are debilitated, the interaction with potentially hazardous environmental factors may result in internal dysregulation, such as abnormal immune activation, increased oxidative stress, as well as genomic instability. It has been established that immune system dysregulation, including infections in early pregnancy, prenatal maternal inflammation, and increased neuronal inflammation, could be contributing factors for the development of ASD (Koufaris & Sismani, 2015). Additionally, studies have also shown that environmental factors have the ability to affect epigenetics. These are mechanisms that biochemically modify DNA or histones affecting gene expression, without changing the DNA sequence (Modabbernia, Velthorst, & Reichenberg, 2017).

Environmental factors associated with ASD that can affect these mechanisms include exposure to heavy metals, low levels of dietary folate (Koufaris & Sismani, 2015), as well as maternal use of valproate, used primarily to treat epilepsy and bipolar disorders (Kalkbrenner, Schmidt & Penlesky, 2014), due to its ability to inhibit histone deacetylase and interference with folic acid metabolism (Modabbernia et al., 2017). Air pollution has also been associated with ASD, with a study reporting elevated risk for ASD in adjusted analyses of the top quartile of exposure to chlorinated solvents, heavy metals, diesel particles and other individual compounds (Lyall, Schmidt, & Hertz-Picciotto, 2014). Moreover, exposure to multiple ubiquitous endocrine disruptors (EDCs), such as phthalates, bisphenol A, polychlorinated biphenyls and polybrominated diphenyl ethers has been associated with autism risk.

Seeing as ASD has an early onset development, it comes as no surprise that a number of studies have focus on maternal lifestyle, with attempts to correlate prenatal factors to ASD development. A study found that by every 10 year increase in maternal and paternal age, the increase of ASD risk in the offspring is that of 18% and 21% respectively. In addition to this, it has also been hypothesized that the increase in ASD risk may be attributed to the accumulation of toxins, such as persistent organic pollutants (PCBs) and EDCs, in the body since it is probable that older parents lived in times prior to chemicals bans and experienced more years of bioaccumulation (Kalkbrenner et al., 2014). An efficient way to measure the relationship between ASD and environmental exposures is, ideally, through large case-control studies, which demand objective information and a direct measure of compounds or metabolites in bio-specimens. However, in Portugal, this information and archived samples are not often accessible for use cohort studies. Additionally, this type of prospective studies are commonly complex, take a long period and require a substantial funding, which may lead to incomplete data and biological sample collection, as well as parents dropping out of the study, In this sense, the inclusive assessment of environmental exposure in ASD in large datasets from variable environmental settings is lacking.

The Early Life Exposure Assessment Tool (ELEAT) was originally developed in California by researchers of University of California Davis. It is a questionnaire that aims to assess environmental exposures that have been associated with ASD development, allowing for an indirect approach given that it uses measurements of exposures in specific microenvironments (e.g. home) and human activity pattern (e.g. maternal lifestyle) data to predict levels of fetal/infant exposure. Indirect exposures, such as this one, are used frequently on account that they require less resources than the assessment of direct exposures. The aim of this study was to pilot a Portuguese version of the Early Life Exposure Assessment Tool (ELEAT) for the assessment of the role of environmental exposures in a population of Portuguese children with ASD.

## METHOD

### Participants

Participants were organized in two groups. The first group was composed of 20 mothers of children with ASD recruited from the Autism Spectrum Disorder in the European Union (ASDEU) project. This project aimed to investigate the prevalence of autism in 12 countries in the European Union, including Portugal, which was finalized in 2017. The second group included 12 co-workers employed by *Instituto Nacional de Saúde Ricardo Jorge* (INSA), whom offspring had no history of neurodevelopmental disorder. These mothers self-administered the ELEAT survey on paper prior to the focus group session.

From the mothers who agreed to participate, all attended the focus group.

### Material

The ELEAT’s objective is to understand how environmental risk factors can influence ASD development.

This tool is divided in 10 modules: Demographic Information, Maternal Conditions/Medical Interventions, Breastfeeding and Child Diet, Maternal Diet, Supplements, Lifestyle, Home and Environment, Environment, and Occupation and Exposures, and Instrument Evaluation. This questionnaire is constituted by 259 items (some may be skipped with gateway questions), corresponding to 80 pages, which intend to asses 3 different periods: it examines maternal exposure to exogenous factors from 3 months before conception to the first year of the child’s life. Therefore, covering crucial phases of early neurodevelopment, with each module analyzing different sources of exposure.

In addition to the paper version, an online version was created in the RedCAP survey, a secure web application for building and managing online surveys and databases. Here the mothers were able to access the questionnaire through a link, and were given the option to save their answers and proceed filling it in later. This became the main filling method not only because it was more organized, but mainly because, seeing as the questionnaire is extensive, the fact that the mothers could save their answers and return to it later made it easier and more appealing due to busy lifestyles. However, the mothers had always the chance to fill out the questionnaire in paper version.

### Procedure

Mothers of children with ASD and typically-developing children filled the Portuguese version of ELEAT, which was translated from the United Kingdom version due to the greater similarities in lifestyle and culture compared to the original United States version. For example, the American version of the tool included questions regarding fish types that are not consumed in Portugal, as well as housing typologies. Cross-cultural problems that might compromise its validity are minimized by European similarities regarding lifestyle, habits, dietary, household and other daily items, comparable access to healthcare and education, as well as employment and infrastructure, and no major cultural adaptations were made.

After being translated, the ELEAT was piloted in a group of mothers of typically-developing and mothers of ASD children. It was mostly completed online, although a paper version was also available, with an average completion time of 60 minutes.

Focus Groups were conducted by the same psychologist with experience in exploring health and environmental themes. Both focus groups were supported by specific scripts, which was addressed towards the improvement of questions and modules, determine the most logical question order, simplify formatting, and shorten the instrument. For this, we focused on the Instrument Evaluation module, which allows the participants to evaluate the length, comprehensiveness, answers certainty and relevance of the tool.

Simultaneously, another research member took notes of salient patterns, answers, suggestions and themes from the discussions.

Both focus group sessions were recorded, and took around 1 hour and half.

## RESULTS

Two focus groups were constituted to obtain a broad perspective with respect to socio-demographic characteristics of the sample (Table 1). Within each focus group the characteristics of the mothers were homogeneous, however, between focus groups these were more heterogeneous, especially regarding the group with mothers of children with ASD. All mothers were, at least, 18 years of age, with an average of 40 years old, although mothers with ASD children, in Group 1, had a broader age range. Mothers had one or more children. Typically-developing children and ASD children’s age ranged from 2 to 12 years. Most mothers, in both groups, were born in Portugal and spoke Portuguese fluently at home (75% vs. 100% respectively). 92% with typically-developing children and 45% of mothers of children with ASD had a university degree.

**Table 1.** Socio-demographic characteristics of the sample

|  | Children with ASD | Typically-developing children |
| --- | --- | --- |
| Child age mean (age range) | 9.3 (8-12) | 5.0 (2-10) |
| Mother age mean (age range) | 39.6 (32-50) | 40.5 (36-45) |
| Born in Portugal % | 75 | 100 |
| Resident in Portugal % | 100 | 100 |
| Education |  |  |
| Basic % | 55 | 8 |
| University degree % | 45 | 92 |

The focus group discussions were productive and produced important adaptations for the improvement of the ELEAT. Despite the heterogeneity between groups, identical opinions and suggestions arose.

Concerning its length, in general, all mothers agreed that the questionnaire was too long and should be shortened. 34% of the mothers considered the ELEAT very long, and 56% considered it slightly long (Figure 1).

**Figure 1.**
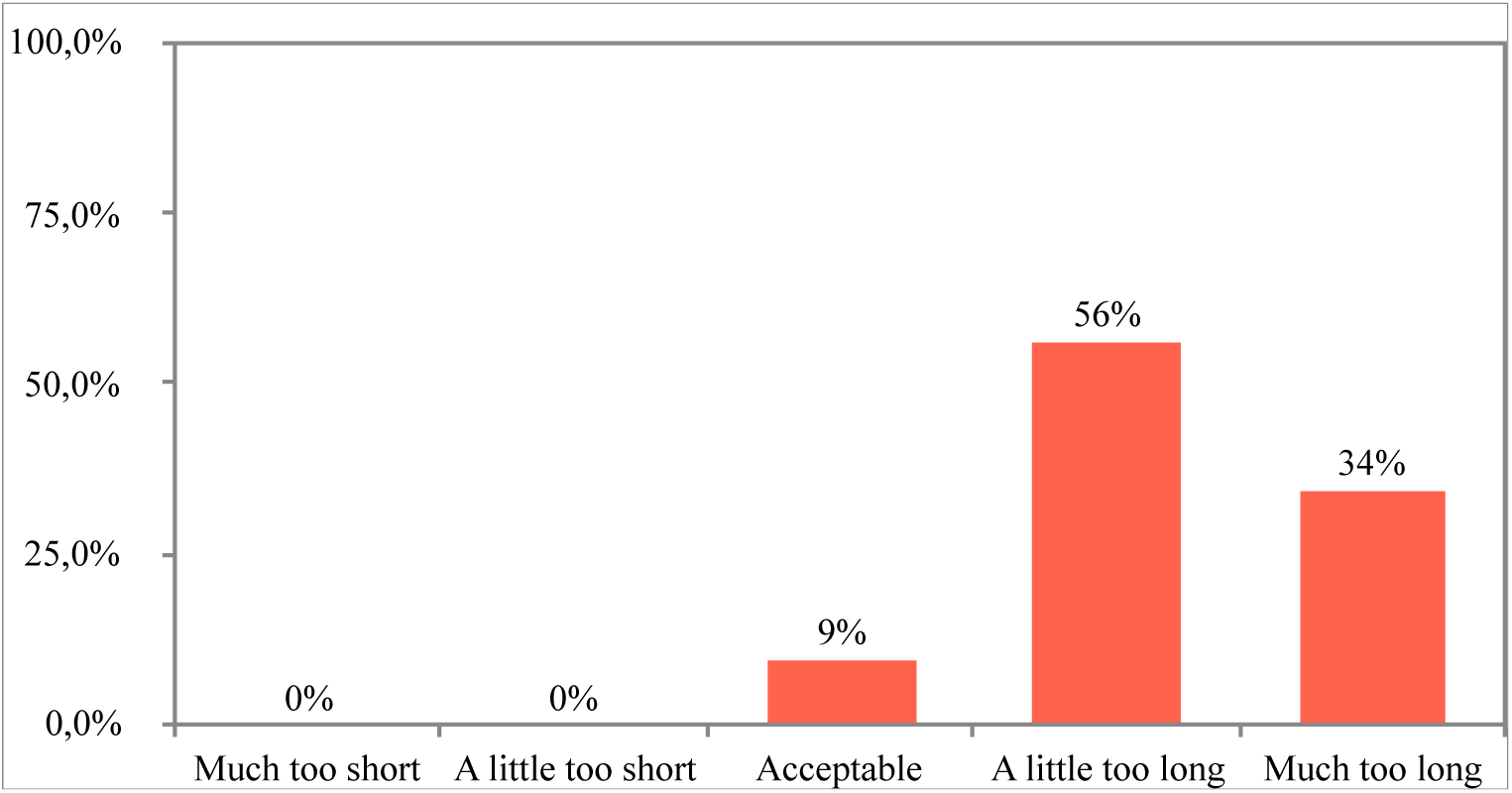
Overall questionnaire length rating (% answers)

Nevertheless, by the end of the discussions these same women felt that it would be best to keep as many questions as possible to avoid omitting potentially important exposure information. In a Lickert-type scale, 63% of mothers agreed and 20% of mothers somewhat agreed that the questions were important, but 10% were neutral (Figure 2).

**Figure 2.**
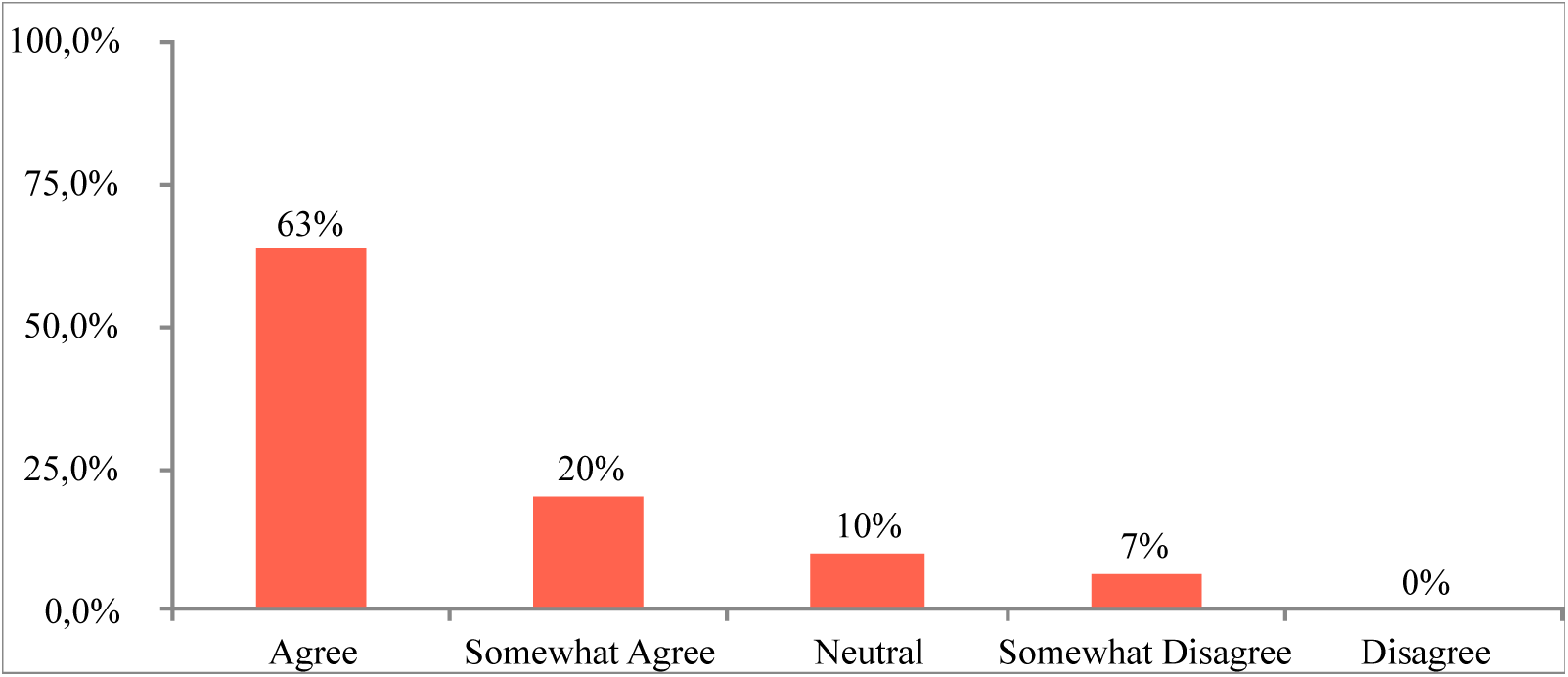
Percentages of importance attributed by the mothers to the tool questions

As Figure 3 shows, 48% agreed and 45% slightly agreed that they were sure about their answers. In fact, one of the concerns discussed and addressed was the certainty level of the given answers, seeing as some mothers gave birth many years ago, and the questionnaire is very comprehensive, requiring them to recall detailed information regarding their pregnancy. However, the strategy adopted by most mothers was to think about their current daily habits, considering that these were not significantly altered during pregnancy.

**Figure 3.**
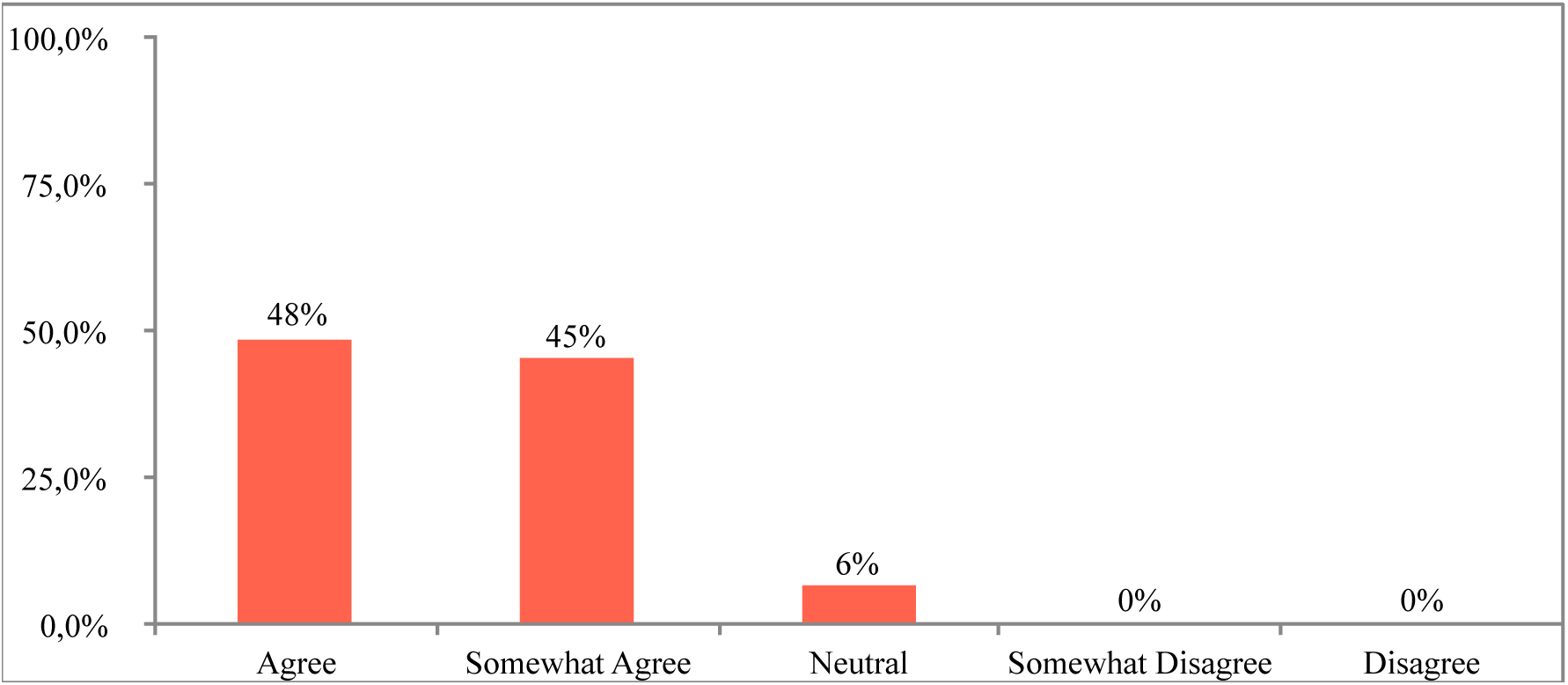
Certainty regarding answers given the mothers for the different modules (% answers)

In all modules, mothers reported above 50% of high confidence about their answers. In four modules (Maternal Condition/Medical History, Breastfeeding, Maternal Diet and Home Environment) between 70% and 88% of the mothers reported high certainty concerning their given answers. Only in Occupation and Exposures did the mothers express lower confidence, with about 52% reporting high certainty (Figure 4).

**Figure 4.**
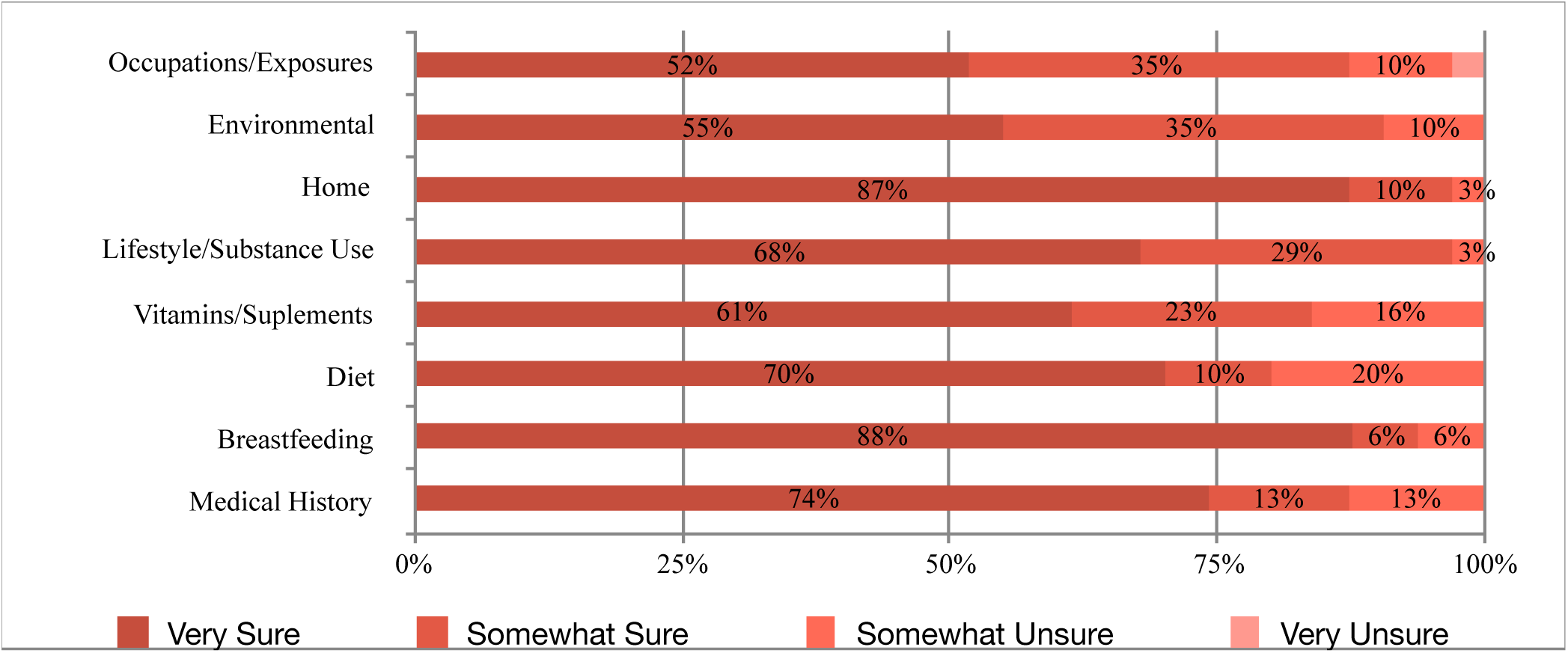
Certainty regarding answers for the different modules (% answers)

**Figure 5.**
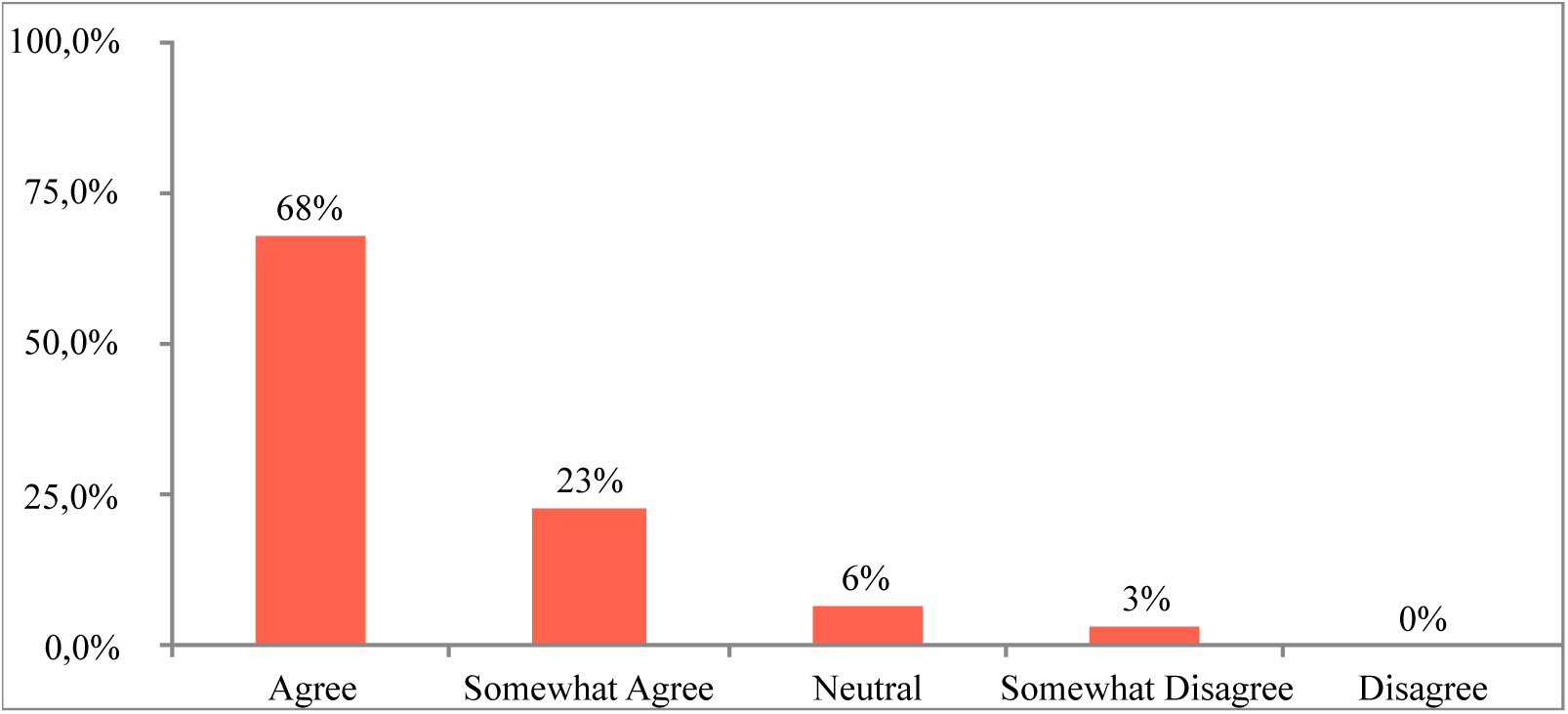
Percentages of clarity about questionnaire questions

The difficulty to recall detailed information from pregnancies was dependent on the subject matter and can be influenced by the interval of time between the pregnancy events and survey administration.

In general, similar concerns were reported in both group discussions regarding the length, detailed and complexity of the questionnaire. The main positive conclusions were that the questions were direct and well written, with 68% of the mothers agreeing and 23% slightly agreeing that the instructions were clear. However, mothers suggested that general vocabulary, specifically medical vocabulary, should be replaced with simpler words and phraseology to broaden survey use for individuals with limited language proficiency.

In general, all mothers favored moving the questionnaire from paper version to online administration, as well as suggesting that the administration should be assisted, thus allowing a more efficient and easier completion. Mothers also favored the flexibility provided by online administration, allowing them to complete the survey in multiple sessions as their schedules allowed.

Another very relevant result was that no question scored 20% regarding the “Do not know” option. In fact, only 3 questions scored 19% on the “Do not know” answer and these questions belonged to the supplements module (Mothers reporting intake of: Vitamin B6?, Vitamin B12?, Calcium?)

## DISCUSSION

The goal of this study was to pilot the Portuguese version of the Early Life Exposure Assessment Tool (ELEAT) for the assessment of environmental exposures roles in a population of Portuguese children with ASD. For this, we conducted two focus groups in order to generate a discussion focused on question clarity, content, module length, formatting, and mode of administration of the tool.

First and foremost, we conclude that both focus group discussions were fundamental to improve and adapt the usability and contents of the ELEAT. In both groups, all mothers participated actively and yielded important suggestions, adaptations, thoughts, ideas, experiences and advices. All content discussed in the focus group sessions was crucial and indispensable for the update of both structural reviews and some vocabulary and details of instrument, making it less dubious and more accessible to broader socio-economic levels of the population.

Morgan (1997) defined focus groups as a research technique that collects data through group interactions about a particular topic suggested by the researcher. It can also be defined as a resource to understand the process of constructing the perceptions, attitudes and social representations of groups of human beings (Smith, 2000).

The starting point for conducting a research project whose feasibility is based by the realization of focus groups is the clarity of its purpose. Methodological decisions depend on the objectives established. This will influence the composition of the groups, the number of elements and the homogeneity or heterogeneity of the participants (culture, age, gender, social status, etc.). All of these factors can influence the discussion process and the finals results. We think that our results were neither influenced nor limited by these factors. Although mothers were not randomly selected from the population, our sample size was relatively small and each group was relatively homogeneous regarding to socio-demographic characteristics. Overall, mothers presented a variety of thoughts, suggestions, ideas and doubts. Most importantly, these perspectives were similar between groups.

The ELEAT instrument was upgraded accordingly to focus groups considerations. Firstly, clinical content was reduced and we believe this could minimize the redundancy and increase user-friendliness so that the mothera can easily self-administer the ELEAT. In addition, this simplification may decrease the literacy level required, and consequently improve data collection accuracy for those with less education. It is very important that the ELEAT is sensitive to cultural factors, embraces diversity and fosters an atmosphere in which the varied life experiences of parents are recognized and respected. Before this pilot study, the ELEAT had already undergone a pilot studies in the US and UK. Our pilot study also allows for the further improvement of this instrument prior to making the survey available for global use. Secondly, results also showed that mothers felt the instructions to answer the tool were clear, and the majority felt sure about their answers. The mothers also appreciated the possibility of filling it online, and gave very positive feedback regarding this process. Feedback from the Instrument Evaluation module is essential, considering that this tool is still being molded to improve its capacity to assemble accurate data from diverse populations. Due to the use of scientific terms in the questionnaire, it is especially important to offer assistance to participants with lower education in order to guarantee accurate data collection. By tackling this and the length of the questionnaire, it is plausible that the selection of “don’t know” option will decrease.

Therefore, ELEAT’s ability to measure multiple factors simultaneously gives it an advantage over studies that measure single exposures. It is important to note that this tool aims simply to gather information about environmental exposure and is not to be used as a diagnosis instrument. It is also important to note that sole exposure to such environmental factors is not sufficient to set in motion the mechanisms that lead to neurodevelopmental problems. It is however the combination of exposure and genetic susceptibility to such factors that may lead to neurodevelopmental consequences.

Taking into account that this is a standardized tool, the ELEAT has the potential of being applied in different countries and gather a large amount of data that could be analyzed together and lead to meaningful findings. It has been noted that the translation of instruments, such as the ELEAT, may present some cross-cultural problems, thus compromising its validity. However, in this pilot study, these are minimized by the fact that there is not a large discrepancy between the lifestyle and habits between British and Portuguese mothers.

The main limitation of this tool was its length. As seen by the results section, this tool was considered to be extensive. This could be one of the principal reasons that could limit the sample size of future studies, given that mothers have to be motivated to fill out the entire questionnaire.

The questions have proven themselves complex to extract quantitative information from, which is necessary for the creation of a score. Therefore, the creation of a rational to transform qualitative into quantitative results is a crucial step to improve the statistical power of ELEAT conclusions.

This pilot study was an important step for future studies that intend to use the ELEAT to identify environmental risk factors for ASD. In fact, there has been an accumulation of evidence pointing to the role played by environmental factors in ASD risk, and, given the difficulty of obtaining information on the environmental exposure, ELEAT can assume a very important role in case– control studies. Currently, research that focus on the role of environmental exposures during fetal development and early postnatal time periods in ASD depend on studies that quantify different analytes and metabolites in biological samples collected from the pregnant mother and/or the child. Additionally, studies that evaluate air quality data regarding prenatal exposures are also performed. Case–control studies based on tools like the ELEAT, which enable the collection of multiple environmental exposure data, are necessary.

The identification of early biological markers requires a specimen collection in firs years. Cohort studies fill all these gaps, and the ELEAT study can therefore provide an excellent tool for prospectively assessing environmental risk and protective factors, collecting biological markers that may correspond to early signs of ASD.

In the future, a combination between the exposures reported in the ELEAT with genetic data collected from the same participants might help reach more solid conclusions. Integrating genetic data could potentially provide a biological explanation to why the potential factors found by the ELEAT may be associated with ASD risk.

## CONCLUSIONS

The main positive conclusions from this piloting of the ELEAT instrument were that the questions were generally considered clear and important, and that most mothers did not have major difficulties recalling details and timing of exposures occurring during their children’s early life. Some concerns were however reported regarding the length and complexity of the questionnaire.

Integration of the pilot feedback will allow us to enhance the tool for its application in Portuguese-speaking communities, improving its capacity to assemble accurate environmental data from diverse cultural settings, and to be extended to larger population datasets. Combined with genetic and clinical data, the ELEAT will contribute to the identification of modifiable lifestyle and environmental risk factors for ASD. Such evidence may eventually provide the opportunity for disease prevention or reduced severity by mitigating exposure when genetic susceptibility is identified early in life.

